# Characterization of flagellar and toxin phase variation in *Clostridium difficile* ribotype 012 isolates

**DOI:** 10.1101/256883

**Authors:** Brandon R. Anjuwon-Foster, Natalia Maldonado-Vazquez, Rita Tamayo

## Abstract

*Clostridium difficile* causes diarrheal diseases mediated in part by the secreted toxins TcdA and TcdB. *C. difficile* produces flagella that also contribute to motility and bacterial adherence to intestinal cells during infection. Flagellum and toxin gene expression are linked via the flagellar alternative sigma factor, SigD. Recently, we identified a “flagellar switch” upstream of the early flagellar biosynthesis operon that mediates phase variation of both flagellum and toxin production in *C. difficile* strain R20291. However, we were unable to detect flagellar switch inversion in *C. difficile* strain 630, a ribotype 012 strain commonly used in research labs, suggesting the strain is phase-locked. To determine whether a “phase locked” flagellar switch is limited to 630 or present more broadly in ribotype 012 strains, we assessed the frequency and phenotypic outcomes of flagellar switch inversion in multiple *C. difficile* ribotype 012 isolates. The laboratory-adapted strain JIR8094, a derivative of strain 630, and six clinical and environmental isolates were all found to be phase OFF, non-motile, and attenuated for toxin production. We isolated low frequency motile derivatives of JIR8094 with partial recovery of motility and toxin production, and found that additional changes in JIR8094 impact these processes. The clinical and environmental isolates varied considerably in the frequency by which flagellar phase ON derivatives arose, and these derivatives showed fully restored motility and toxin production. Taken together, these results demonstrate heterogeneity in flagellar and toxin phase variation among *C. difficile* ribotype 012 strains, and perhaps other ribotypes, which could impact disease progression and diagnosis.

**Importance:** *Clostridium difficile* causes diarrheal disease resulting in significant morbidity and mortality in many countries. *C. difficile* produces flagella that enhance bacterial motility, and secretes toxins that promote diarrheal disease symptoms. Previously, we found that production of flagella and toxins are co-regulated via a flippable DNA element termed the “flagellar switch”, which mediates the phase variable production of these factors. *C. difficile* can exist as flagellated, motile, toxigenic bacteria (“*flg* ON”) or aflagellate, non-motile, nontoxigenic bacteria (“*flg* OFF”) due in part to the flagellar switch orientation. Here we evaluate multiple isolates of *C. difficile* ribotype 012 strains and find them to be primarily *flg* OFF. Some, but not all, of these isolates showed the ability to switch between *flg* ON and OFF states. These findings suggest heterogeneity in the ability of *C. difficile* ribotype 012 strains to phase vary flagellum and toxin production, which may broadly apply to pathogenic *C. difficile*.

## Introduction

The obligate anaerobe *Clostridium difficile* is a leading cause of nosocomial intestinal infections. *C. difficile* associated infections (CDI) are most common in individuals who have undergone antibiotic therapy, which disrupts the usually protective microbiota and creates a niche for *C. difficile* outgrowth (1). Virulence of *C. difficile* is largely mediated by two glucosylating toxins, TcdA and TcdB, which target and inactivate Rho and Rac GTPases in the intestinal epithelium, resulting in depolymerization of the actin cytoskeleton and eventually host cell death (2). Toxin-mediated damage to the epithelium leads to disruption of the intestinal barrier, diarrheal symptoms, and a robust inflammatory response (2).

*C. difficile* produces peritrichous flagella that are essential for swimming motility and contribute to host cell adherence (3-6). As in other bacterial species, the expression of flagellar genes occurs in a hierarchical manner (7). In *C. difficile*, at least four different operons encode flagellar genes. The early stage flagellar genes *(flgB* operon) are transcribed first, and they encode the basal body, motor, and rod of the flagella. The *flgB* operon also encodes the alternative sigma factor SigD (σ^D^, also known as FliA or σ^28^). In addition to activating late stage flagellar gene expression, SigD also positively regulates the expression of *tcdR*, which encodes a sigma factor that activates transcription of *tcdA* and *tcdB* (7-9). Therefore, the regulation of flagellar genes also impacts virulence by affecting toxin gene expression.

The expression of flagellum and toxin genes is subject to complex regulation (6, 7, 10-12). Recently, we demonstrated that flagellum and toxin biosynthesis is phase variable via site-specific recombination that inverts a DNA element termed the “flagellar switch” (13). The flagellar switch consists of a 154 bp invertible DNA sequence flanked by 21 bp inverted repeats and lies upstream of the *flgB* operon. The orientation of the flagellar switch controls expression of the *flgB* operon, and therefore *sigD* and the toxin genes, through an unidentified mechanism occurring post-transcription initiation (13). Bacteria with the flagellar switch in an orientation resulting in flagellum production, swimming motility, and high toxin production were termed flagellar phase ON (*flg* ON). In contrast, bacteria with the flagellar switch in the opposite orientation resulting in the absence of flagella, sessility, and reduced toxin production were termed flagellar phase OFF (*flg* OFF). RecV, a site-specific tyrosine recombinase, mediates inversion of the flagellar switch (13). The phase variable production of flagella and toxins was proposed to allow *C. difficile* to balance the benefits of swimming motility and toxinogenesis with the cost of producing these immunogenic factors (14-17).

Flagellum and toxin phase variation in *Clostridium difficile* appears to vary across ribotypes (13). For *C. difficile* strains R20291, a ribotype 027 strain, and ATCC 43598, a ribotype 017 strain, both the *flg* ON and OFF orientations were apparent under multiple conditions tested. However, only the *flg* ON orientation was detectable in 630, a ribotype 012 strain originally isolated from a patient with *C. difficile* infection (18). However, recent work from Collery *et al.* suggests the flagellar switch is capable of inversion in 630 (19), suggesting this strain is phase-locked. JIR8094 (also named 630E) was isolated through serial passage of 630 to obtain erythromycin-sensitive isolates amenable to genetic manipulation with tools relying on an erythromycin resistance cassette (20). Comparison of the genome sequence of JIR8094 to 630 revealed numerous secondary mutations that arose during and since isolation (19). One polymorphism identified was an inversion of the flagellar switch to the OFF orientation in JIR8094 compared to 630, which is *flg* ON. The JIR8094 genotype explains the non-motile phenotype and reduced flagellum and toxin gene expression previously reported for JIR8094 (7), and indicates an ability to invert the flagellar switch in this strain lineage. In contrast, the 630Δ*erm* strain was similarly derived from 630 through serial passaging, but it remains motile and toxinogenic (21).

In this study, we examined multiple ribotype 012 strains, including laboratory-adapted, clinical, and environmental isolates, to assess their ability to invert the flagellar switch and to phase vary flagellum and toxin production. Analysis of these strains and their motile derivatives indicates that flagellum and toxin phase variation is conserved in ribotype 012 *C. difficile*, and that there is considerable variation in the frequency of phase variation in this and perhaps other ribotypes.

## Materials and Methods

### Bacterial strains and growth conditions

All bacterial strains used in this study are listed in Table S1. *C. difficile* strains were cultivated statically at 37°C in brain heart infusion medium (BD, Beckton Dickinson) supplemented with 5% yeast extract (BHIS) or tryptone yeast (TY) medium, as specified. All *C. difficile* growth was done anaerobically in a Coy anaerobic chamber with an atmosphere of (90% N2, 5% CO2, 5% H2). Unless otherwise indicated, *E. coli* was cultured at 37°C under aerobic conditions in Luria Bertani (LB) medium. For selection of plasmids in *E. coli*, 100 μg/mL ampicillin (Amp) and/or 10 μg/mL chloramphenicol (Cm) was used, as indicated. Kanamycin (Kan) 100 μg/ml was used to select against *E. coli* in conjugations. For maintenance of plasmids in *C. difficile*, 10 μg/mL thiamphenicol (Tm) was used.

### Characterization of *C. difficile* ribotype 012 clinical and environmental isolates

Clinical and environmental isolates of *C. difficile* used are part of the F ribotype database (22). The strains were ribotyped by fluorescent PCR and sequencing of ribosomal RNA genes as previously described (22). The flagellar switch orientation was determined by PCR amplification of the region and sequencing of the products using primers R591 and R857.

### Orientation-specific PCR

The flagellar switch orientation was determined by PCR using two primer sets that distinguish the orientation of the switch (13, 23). Primers were designed using the published genomes sequences for strains 630 (Accession No. FN688375.1) and R20291 (Accession No. FN545816.1). Primer sequences are listed in Table S2. For the 630 lineage and other ribotype 012 isolates, R1751 and R857 were used to detect the ON orientation, and R1752 and R857 were used to detect the OFF. For R20291, R1614 and R857 were used to detect the ON orientation, and R1615 and R857 were used to detect the OFF. PCR products were separated on a 2% agarose gel and stained with ethidium bromide (EtBr) for imaging using the G:Box Chemi imaging system.

### Motility assays

Swimming motility was assayed as described previously (13, 24). Briefly, individual colonies of the indicated strains were grown in TY broth overnight, then diluted 1:50 in fresh BHIS and grown to an OD_600_ ~ 1.0. In standard assays, 2 μL of culture were inoculated into anaerobic motility medium (0.5X BHIS-0.3% agar, autoclaved for 30 minutes). In experiments aimed at identifying JIR8094 motile revertants, a range of volumes of overnight bacterial cultures were centrifuged to collect and concentrate bacterial cells. The cells were suspended in 100 μL of phosphate buffered saline (PBS), and 2 μL were stabbed into soft agar plates. Swimming motility was assayed by measuring the colony diameter after 24, 48, and 72 hours incubation as indicated. Two perpendicular measurements were averaged for each colony. Photographs shown were taken with the G:Box imaging system (Syngene). To measure the frequency of motile recovery in the ribotype 012 isolates, motility assays were performed as described above with 32 independent colonies in two independent experiments. A stab site positive for motility was identified by the presence of a motile flare.

### Recovery of motile derivatives of parent JIR8094 and select ribotype 012 isolates

After incubation in motility agar as described above, motile bacteria were collected from the edge of an expanded colony using a sterile loop and passaged onto BHIS agar for recovery. All isolates were confirmed to be motile by independently assaying swimming through motility medium, and the orientation of the flagellar switch was determined by PCR as described.

### Microscopy

For transmission electron microscopy, colonies of *C. difficile* were grown in TY medium overnight, diluted into fresh BHIS medium, and grown to an OD600 of ~1.0. Bacteria were washed in Dulbecco’s PBS (Sigma) prior to suspension in PBS-4 % paraformaldehyde for fixation for 1 hour in the anaerobic chamber. Cell suspensions were adsorbed onto Formvar/copper grids, washed with water, and stained for 30 seconds in two sequential drops of aqueous uranyl acetate (1.5%). Cells were viewed on a LEO EM 910 Transmission Electron Microscope (Carl Zeiss Microscopy, LLC, Thornword, NY). Documentation was done with a Gatan Orius SC1000 digital camera and Digital Micrograph 3.11.0 software (Gatan, Inc., Pleaston, CA).

### Detection of TcdA production by western blot

Overnight cultures were grown in TY broth and then diluted 1:50 in fresh medium. Strains were grown until late stationary phase (OD_600_ 1.8-2.0) for 24 hours, normalized to the same density, and then cells were collected by centrifugation. Bacteria were lysed by suspension in SDS-PAGE buffer and heating to 100°C for 10 minutes. Samples were electrophoresed on 4-15 % TGX gels (Bio-Rad), then transferred to nitrocellulose membranes. Membranes were stained with Ponceau S (Sigma) to determine equal loading and imaged using the G:Box Chemi imaging system. TcdA was detected using mouse anti-TcdA primary antibodies (Novus Biologicals) followed by goat anti-mouse IgG secondary antibody conjugated to DyLight 800 4X PEG (Invitrogen). Blots were imaged using the Odyssey imaging system (LI-COR). All strains were assayed in at least 3 independent experiments.

### RNA extraction, cDNA synthesis and quantitative reverse transcriptase–PCR

*C. difficile* was grown in TY medium overnight, diluted in fresh medium, and grown until mid-exponential phase (OD600 ~1.0) or stationary phase (OD600 1.8-2.0) as indicated. Cells were collected by centrifugation, and RNA was isolated as described previously (13, 24). RNA was treated with DNase I (Ambion), and cDNA was then synthesized using the High-Capacity cDNA Reverse Transcription Kit (Thermo Fisher). qRT-PCR was done using SensiMix SYBR and Fluorescein Kit reagents (Bioline). The primers used were named following the convention gene-qF and gene-qR (Table S2). Data were normalized using the ΔΔC*t* method, with *rpoC* serving as the reference gene. At least four independent samples were included.

### Generation of *topA* expression strains

The *topA* gene was amplified from 630 genomic DNA using primers R2302 and R2303, which introduced SacI and BamHI restriction sites, respectively. After digestion with these enzymes, the *topA* fragment was ligated into similarly digested pRPF185, which allows for ATc-inducible expression of the cloned gene (25). Ligation products were transformed into *E. coli* DH5α. Chloramphenicol-resistant clones were collected and screened by PCR with R2302 and R2303. Clones were confirmed by sequencing of the insert. The resulting pRPF185∷*topA* was introduced into *C. difficile* 630, JIR8094, and the motile derivatives of JIR8094 by conjugation via HB101(pRK24) as previously described (24, 26, 27). Transconjugants were selected on agar media with Tm and Kan and confirmed by PCR with plasmid specific primers R1832 and R1833. To induce expression of *topA*, anhydrotetracycline was added at a final concentration of 20 ng/mL to motility medium to assay swimming motility and broth medium for assaying TcdA production by western blot.

## Results

### The laboratory-adapted *C. difficile* strain JIR8094 is *flg* OFF

The *C. difficile* strain JIR8094 is a non-motile, toxin-attenuated derivative of the ribotype 012 strain 630 (7, 19, 20). In *C. difficile*, expression of the *flgB* operon is controlled by multiple mechanisms, including a σ^A^-dependent promoter (28), a c-di-GMP riboswitch in the 5’UTR (24, 28, 29), and the flagellar switch (13). Sequencing of the *flgB* operon regulatory region from *C. difficile* 630 and JIR8094 revealed no differences in the promoter region or the c-di-GMP riboswitch sequences. However, while the LIR and RIR were identical in the two strains, the intervening 154 bp was inverted in JIR8094 relative to 630 (Figure 1A). Alignment of the flagellar switch sequence using the reverse-complement of the sequence obtained for JIR8094 restored sequence identity to 100% with 630 (Figure 1A). Inversion of the flagellar switch in JIR8094 relative to 630 was confirmed by orientation-specific PCR (Figure 1B). For this assay, a common reverse primer (Rv) is used in combination with a primer (Fw1) that leads to amplification when the template is in the ON orientation, or with a primer (Fw2) that amplifies when the template is in the OFF orientation (Figure 1B) (13). As shown previously, *C. difficile* R20291 cultures contain a mixture of *flg* ON and OFF bacteria (Figure 1C) (13). In contrast, only the *flg* ON orientation was detected for strain 630, and only the *flg* OFF orientation was detected for JIR8094 (Figure 1C). In addition, strain 630 demonstrated swimming through motility medium, while JIR8094 was non-motile after 72 hours (Figure 1D). These observations are consistent with a recent report identifying the sequence inversion in JIR8094 (19), and with phenotypes corresponding to the *flg* OFF state (7).

**Figure 1.**
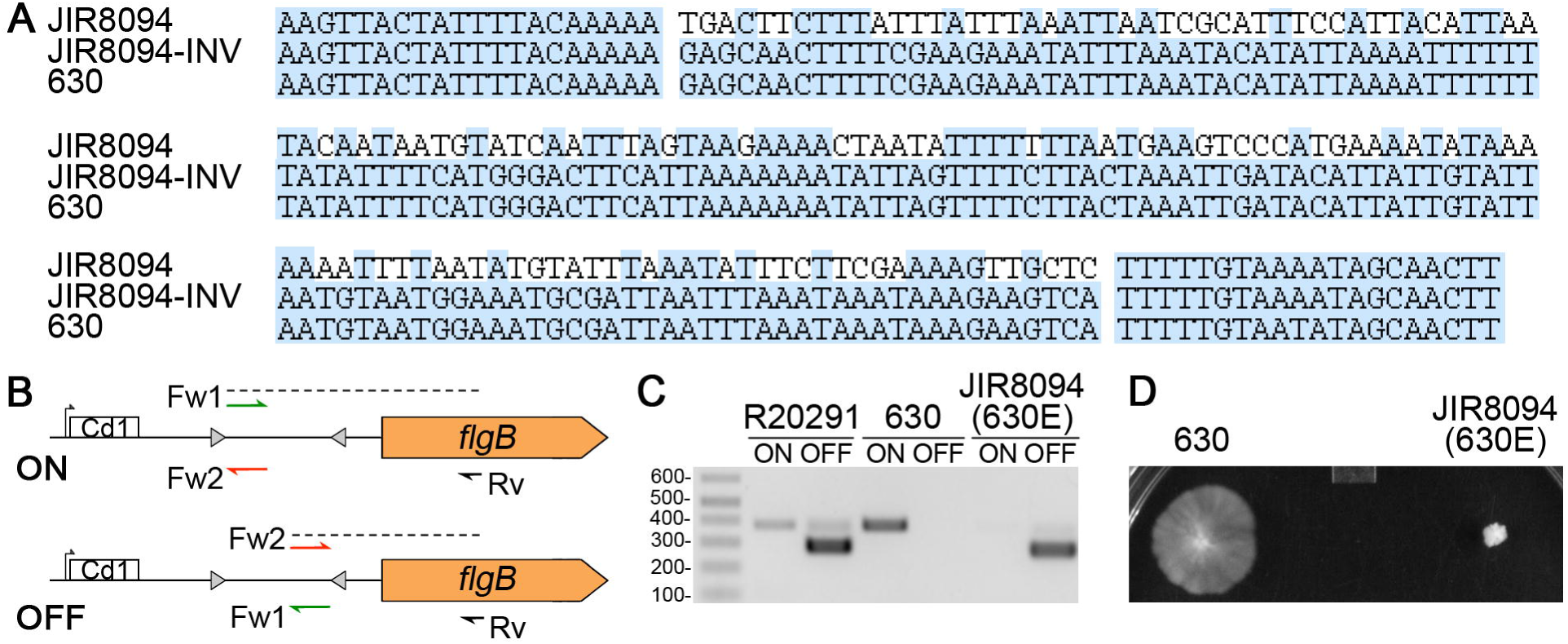
*C. difficile* JIR8094 has the flagellar switch in the OFF orientation. (A) Multiple sequence alignment of the flagellar switch sequences from *C. difficile* 630, its JIR8094 derivative, and JIR8094-inv in which the 154 bp sequence between the imperfect inverted repeats was replaced with the reverse-complement. Blue shading indicates identical nucleotides. (B) Schematic for the orientation specific PCR assay used to determine the orientation of the flagellar switch. The Rv primer (R857) is used with either Fw1/ON (R1751) or Fw2/OFF (R1752) to detect the ON or OFF orientation of the flagellar switch, respectively. In the 630 lineage, Fw1/ON + Rv yield a 374 bp product; Fw2/OFF + Rv yield a 250 bp product. Diagram not drawn to scale. (C) Orientation-specific PCR products using template DNA from *C. difficile* R20291, 630 and JIR8094. (D) Motility of 630 and JIR8094 in motility medium (0.5X BHIS-0.3% agar) after 72 hours growth.

### The flagellar switch undergoes inversion at low frequency in JIR8094

Throughout our experimentation, we did not observe motility or motile flares from JIR8094 in standard assays (data not shown), and Collery et al. similarly reported an inability to recover motile bacteria from this strain (19). These findings suggested that JIR8094 is phase-locked OFF or undergoes inversion of the flagellar switch at a low rate. To distinguish between these possibilities, we modified the motility assay to accommodate higher density inoculums. Rather than inoculating motility agar with 2 μL of saturated broth culture, we first concentrated the culture 30-fold, reasoning that this would increase the odds of detecting rare *flg* ON bacteria. Over 48-72 hours, we were able to detect motility from a subset of JIR8094 inoculums (Figure 2A).

**Figure 2.**
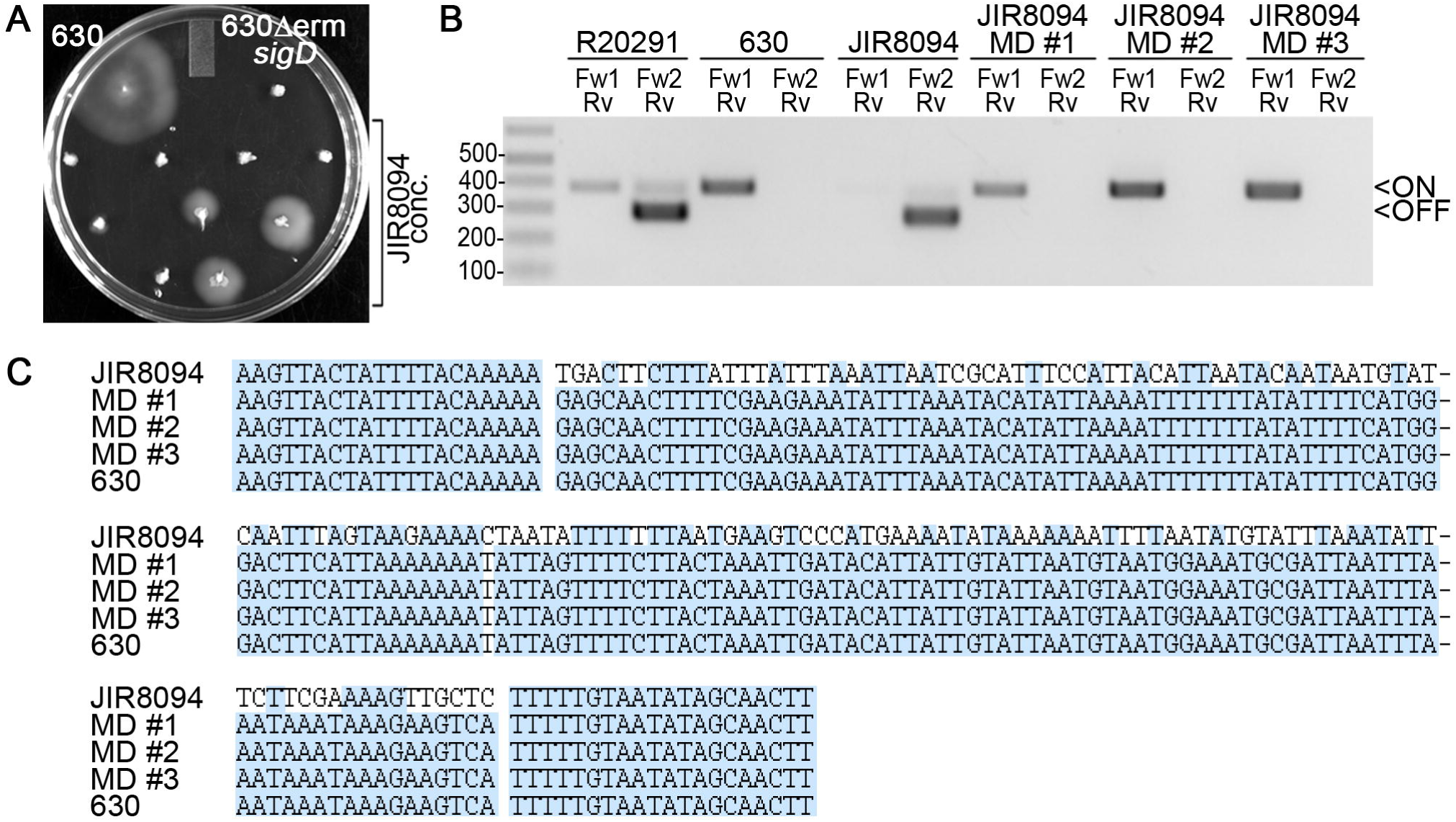
The flagellar switch undergoes inversion at a low frequency in JIR8094. (A) Concentrated liquid cultures of JIR8094 were inoculated into motility medium along with standard (i.e., not concentrated) inoculums of 630 (motile) and 630Δ*erm sigD* (non-motile) as controls. Shown is a representative experiment in which a JIR8094 culture was concentrated 30-fold (9 replicates). After 48 hours growth at 37°C, motility or motile flares was detected for three concentrated inoculums with JIR8094. Data is representative of 3 independent experiments. (B) Orientation-specific PCR products for three motile *flg* ON derivatives of JIR8094 (MD #1 −3). *C. difficile* R20291, 630, and JIR8094 were included as controls for both PCR primer sets. (C) Multiple sequence alignment of the flagellar switch of JIR8094, 630, and three *flg* ON derivatives of JIR8094. Blue shading indicates identical nucleotides.

To determine whether motility was restored as a result of flagellar switch inversion to ON or due to extragenic suppressor mutations, we used orientation-specific PCR (Figure 1B) to assess the orientation of the flagellar switch. Three motile derivatives of JIR8094 (MD #1-3) were arbitrarily chosen after recovery from the edge of the motile colony for further analysis. All three MDs yielded PCR products indicative of the *flg* ON orientation, but not the *flg* OFF orientation, as seen in the parental strain 630 (Figure 2B). Sequencing of the flagellar switch confirmed the *flg* ON orientation of the switch in JIR8094 MD #1-3 (Figure 2C). These results indicate that the flagellar switch remains reversible in JIR8094, though the frequency of inversion appears to be lower than previously observed for R20291 (13).

To estimate the frequency of *flg* ON bacteria in JIR8094, a range of bacterial densities were assayed for swimming motility. As controls, we used the motile 630 strain, the non-motile 630Δ*erm sigD* mutant (30), and an isolate of *C. difficile* R20291 *flg* OFF which contains ~3 % *flg* ON bacteria (13). This small population of *flg* ON bacteria spatially segregates from non-motile *flg* OFF bacteria in motility medium, leading to a motile phenotype that is readily apparent within 24 hours (13). Whereas R20291 appeared motile at all inoculums tested, motility of JIR8094 was only observed at the highest density, suggesting that flagellar switch inversion occurs at a low frequency in this strain (Figure S1).

### Characterization of JIR8094 motile derivatives

We predicted that restoration of the flagellar switch to the ON orientation would restore flagellar motility and toxin production of the JIR8094 MDs to levels observed in the original 630 parental strain. To test this, we assayed swimming motility using standard (not concentrated) inoculums, with *C. difficile* 630 and JIR8094 strains included as positive and negative controls, respectively. The JIR8094 MD #1-3 were motile at all time points tested, unlike JIR8094 (Figure 3A), but motility was not fully restored to the level seen for 630 (Figure 3B). We also observed that the JIR8094 MD #1 produced fewer flagella than strain 630, which displayed many peritrichous flagella; JIR8094 remained aflagellate (Figure 3C).

**Figure 3.**
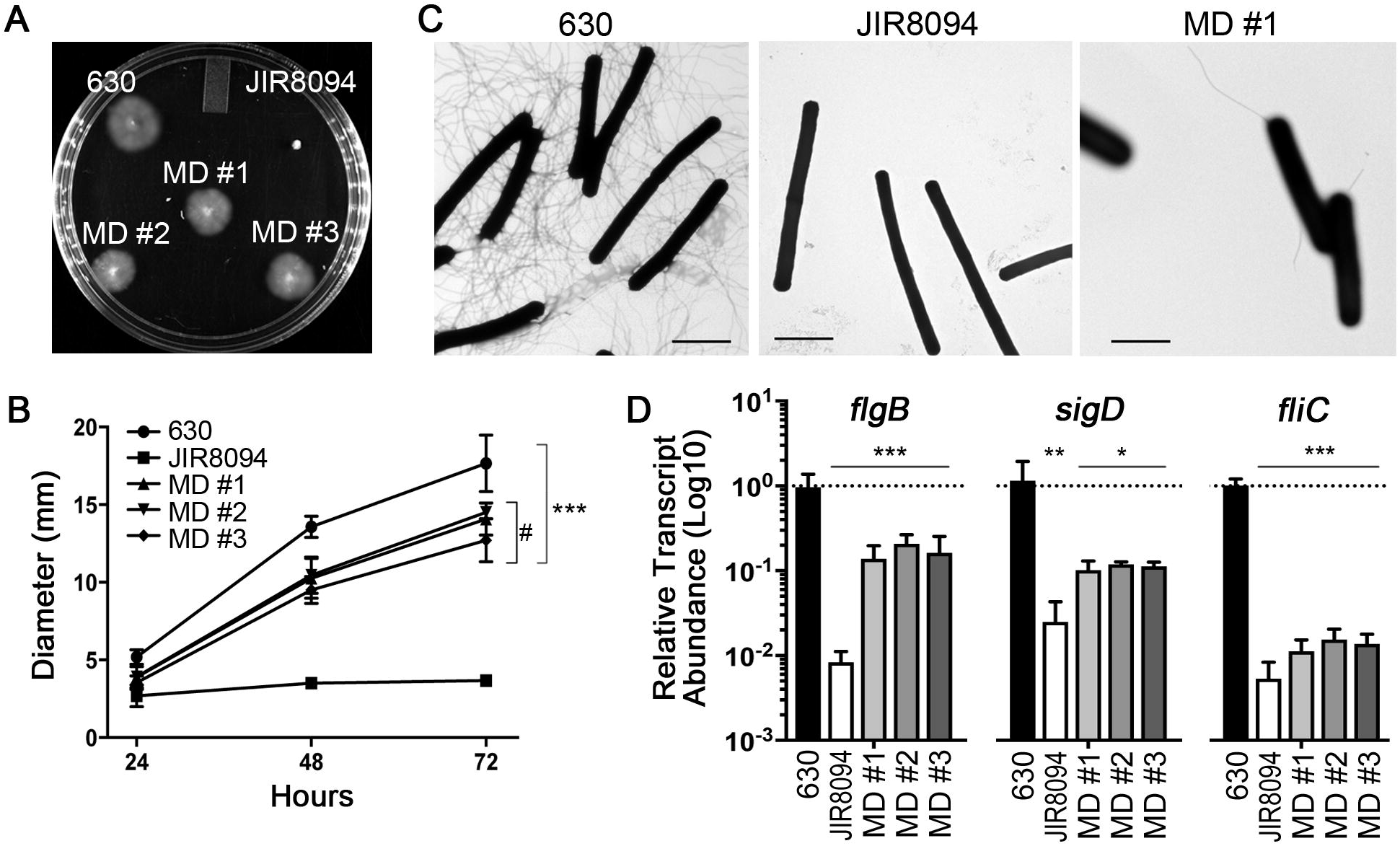
JIR8094 motile derivatives have partial recovery of flagellar gene expression, flagellum biosynthesis, and motility. (A) Motility assay for 630, JIR8094, and JIR8094 MD #1-3. Image is representative of 3 biological replicates. (B) Quantification of motility after 24, 48, and 72 hours. Shown are the means and standard deviations of the swim diameters from 3 independent experiments. At both 48 and 72 hours, *** p < 0.001 compared to JIR8094, and *#* p < 0.05 compared to 630, by one way ANOVA and Tukey’s multiple comparisons test. The motilities of the JIR8094 MD #1-3 were not significantly different from each other. (C) Transmission electron microscopy (TEM) images for 630, JIR8094, and JIR8094 MD #1 showing the presence or absence of flagella. (D) The abundance of flagellar gene transcripts *flgB, sigD*, and *fliC* in logarithmic phase BHIS cultures of 630, JIR8094, and JIR8094 MD #1-3 isolates. The data were analyzed using the ΔΔC*t* method with *rpoC* as the control gene and 630 as the reference strain. Shown are the means and standard deviations. * p < 0.05, ** p < 0.01, *** p < 0.001 compared to the 630 reference by one-way ANOVA and Tukey’s multiple comparisons test. None of the differences between JIR8094 and MD #1-3 values met p < 0.05.

The intermediate motility of JIR8094 MD #1-3 could be due to either partial recovery of flagellar gene transcription and/or of flagellum biosynthesis, both of which could be due to additional mutations in JIR8094. Using quantitative reverse transcriptase PCR (qRT-PCR), we measured the transcript abundance for three flagellar genes, *flgB*, *sigD*, and *fliC*. Both *flgB* and *sigD* are in the operon directly controlled by the flagellar switch (early stage flagellar genes), and *fliC* serves as a representative *sigD*-regulated late stage flagellar gene. As previously reported, the *flgB*, *sigD*, and *fliC* transcripts were significantly lower in JIR8094 compared to 630 (Figure 3D) (7). In MD #1-3, *flgB*, *sigD*, and *fliC* transcript levels were higher than in JIR8094, but were not restored to the levels in 630 (Figure 3D). Nonetheless, the intermediate level of flagellar gene expression and number of flagella on the JIR8094 MDs were sufficient to confer some motility.

Similar intermediate phenotypes were observed with regards to toxin production. Analysis of *tcdR*, *tcdA*, and *tcdB* transcripts in stationary phase cultures (favoring toxin gene expression) by qRT-PCR showed that all three transcripts were less abundant in JIR8094 than in 630, as seen previously (7), although only *tcdA* was significantly lower (Figure 4A). The transcripts trended higher in MD #1-3 than in their JIR8094 parent, but the differences were not statistically significant (Figure 4A). Consistent with this, the amounts of TcdA protein produced by MD #1-3 appeared more similar to the JIR8094 parent than 630 (Figure 4B). Together, these data indicate that JIR8094 MD #1-3 have restored the ability to transcribe flagellar and toxin genes, and thus motility and toxinogenesis, but suggest that other mutations in JIR8094 also affect flagellum and toxin gene expression in this strain.

**Figure 4.**
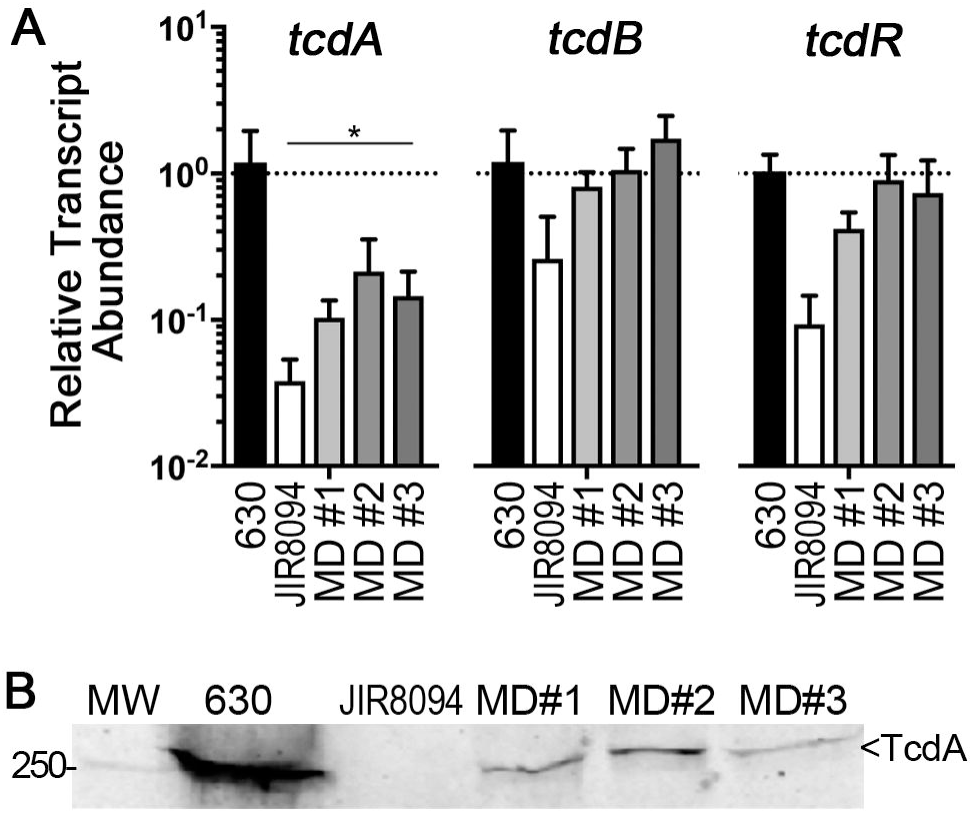
JIR8094 motile derivatives show intermediate toxin gene expression and toxin production. (A) qRT-PCR analysis of *tcdR, tcdA*, and *tcdB* transcript abundance in stationary phase cultures of 630, JIR8094, and JIR8094 MD #1-3. The data were analyzed using the ΔΔC*t* method with *rpoC* as the control gene and 630 as the reference strain. Shown are the means and standard deviations. * p < 0.05 compared to 630 by one-way ANOVA and Tukey’s multiple comparisons test. None of the differences between JIR8094 and MD #1-3 values met p < 0.05. (B) Western blot detection of TcdA production by 630, JIR8094, and JIR8094 MD #1-3. A representative image from 3 independent experiments is shown.

### Topoisomerase activity in JIR8094 motile derivatives impacts motility

Comparison of the genome sequences of *C. difficile* 630 and its 630Δ*erm* and JIR8094 derivatives revealed multiple single nucleotide polymorphisms, including 11 found in JIR8094 but not the other two strains (one of which is the flagellar switch inversion) (19). We speculated that one or more of these polymorphisms is responsible for the difference in flagellar gene expression between 630 and the JIR8094 MD #1-3 (all of which are *flg* ON). Of interest was a nonsense mutation in a *topA* orthologue that introduced a stop codon at amino acid 386, while the full-length protein is 695 amino acids. Because *topA* encodes DNA topoisomerase I, which controls DNA negative supercoiling, we postulated that the mutation in *topA* could result in changes to the *flgB* operon promoter and/or coding sequence that impair gene transcription in JIR8094 and its derivatives, compared to 630 which encodes the full length TopA. If so, expressing TopA in the JIR8094 MD #1-3 is expected restore motility and toxin production to the levels in 630.

To test this, we ectopically expressed *topA* from an ATc-inducible promoter in 630, JIR8094, and the MD #1-3 and compared the motility of these strains to vector controls. A *630Δerm sigD* mutant bearing vector or the *topA* expression plasmid (pTopA) was included as a non-motile control. After 72 hours, expression of *topA* did not significantly alter the motility of 630 or the *sigD* mutant as expected (Figure 5A and 5B). Expression of *topA* alone is not sufficient to augment motility in *flg* OFF bacteria, as JIR8094 + pTopA remained non-motile (Figure 5A and 5B). However, expression of *topA* in JIR8094 MD #1-3 significantly enhanced motility, restoring it to 630 levels (Figures 5A and 5B). Presumably TcdA levels would concomitantly increase upon restoration of TopA activity, but western blot analysis indicated TcdA levels were unchanged in the JIR8094 MD #1-3 + pTopA derivatives compared to parent JIR8094 + pTopA (Figure 5C). These results suggest that while the flagellar switch must be in the ON orientation to allow flagellar gene expression and motility, and DNA supercoiling affects flagellar gene expression. Moreover, additional mutations in JIR8094 may interfere with the link between toxin and flagellar gene expression.

**Figure 5.**
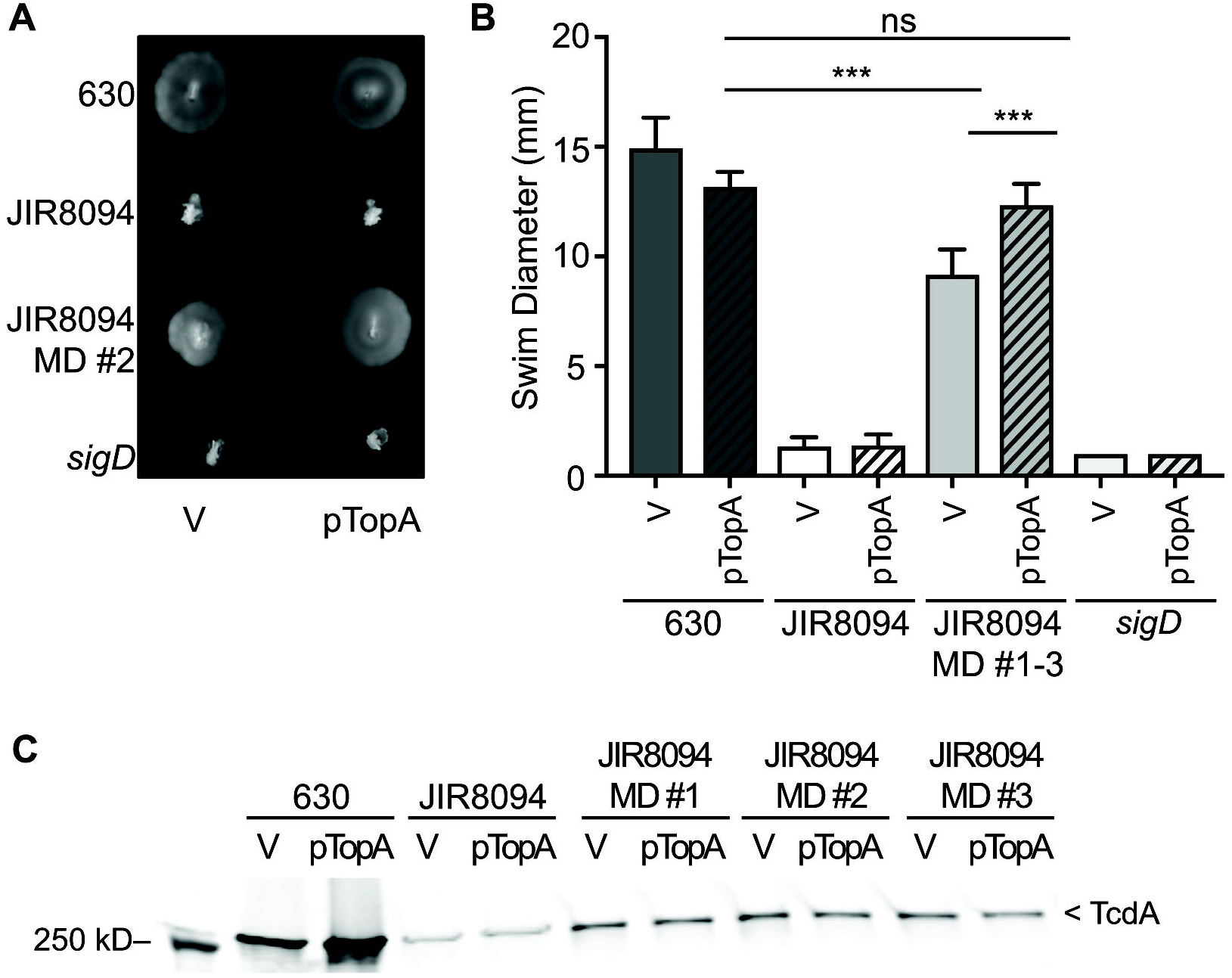
Ectopic expression of *topA* restores motility of JIR8094 motile derivatives to 630 levels. (A, B) Motility of 630, JIR8094, JIR8094 MD #2, and a *sigD* mutant, each bearing vector (V) or pTopA, after 72 hours growth. (A) Image is representative of two independent experiments with six biological replicates of each strain. Equivalent results were obtained for JIR8094 MD #1 and MD #2. (B) Quantification of motility in the above strains after 72 hours, with the data for JIR8094 MD #1-3 combined. Two independent experiments with six biological replicates were done. *** p < 0.001 by one-way ANOVA and Tukey’s multiple comparisons test. (C) Detection of TcdA in the indicated strains by western blot. Data are representative of two independent experiments with each strain.

### Clinical and environmental Isolates of *C. difficile* ribotype 012 are *flg* OFF

Both flagellar switch orientations were readily detected in *C. difficile* strains R20291, a ribotype 027, and ATCC 43598, a ribotype 017, but only the *flg* ON was detected in 630 (13). The 630 strain was isolated in 1982, so passaging within and between labs could have led to accumulation of mutations that reduced the frequency of flagellar phase variation in 630. We thus asked whether the lack of (or very low frequency of) flagellar switch inversion in 630 is unique to this strain and its derivatives, or a broader attribute of currently circulating ribotype 012 *C. difficile* isolates (18). To test this, we evaluated flagellar phase variation in six riboype 012 isolates: SS235, a community environmental isolate; SE838, a hospital environmental isolate; and MAM30, MT4768, MT5065, and MT5066, isolates from patients with CDI (kindly gifted by Dr. Kevin Garey). Sequencing of the flagellar switches revealed that all six isolates contain the flagellar switch in the same OFF orientation as JIR8094, and opposite that of strain 630 (Figure S2) (19). The flagellar switches of two isolates also contained polymorphisms within the inverted repeats (Figure S2). MT4768 had one nucleotide (nt) substitution in the left inverted repeat (LIR) and an additional nucleotide in the right inverted repeat (RIR). SE838 contained an additional 3 nt in the LIR and 2 nt in the RIR. In addition, MT4768 possessed three single nucleotide substitutions within the flagellar switch (Figure S2). SS235, MAM30, MT5065, and MT5066 had flagellar switches identical to that in JIR8094.

Sequencing results indicate the conservation of the regulatory flagellar switch element, but do not reveal if it is functional. It is also possible that the sequencing results reflect the most abundant switch orientation in a population, masking the presence of a lower abundance orientation. Therefore, we used PCR with orientation-specific primer sets that discriminate between the two DNA orientations (Figure 6A) (13). As a control we included 630, for which only the *flg* ON orientation is detectable (Figure 6A) (13). For all six isolates only the *flg* OFF orientation of the flagellar switch was detectable (Figure 6A).

**Figure 6.**
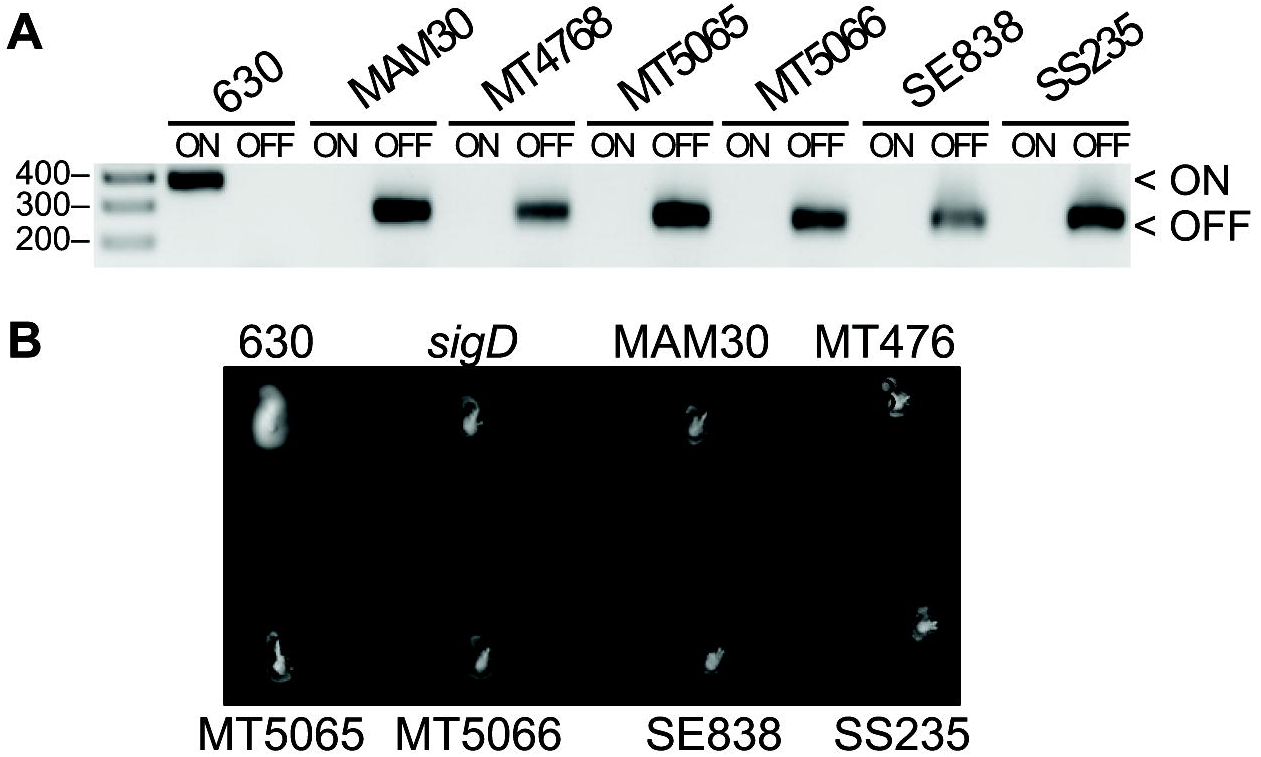
Clinical and environmental 012 ribotype isolates are *flg* OFF. (A) Orientation-specific PCR products using the indicated strains as template. Template from 630 is included as a control to detect the *flg* ON orientation. Image is representative of four biological replicates. (B) Swimming motility after 24 hours. Data are representative of three independent experiments with eight biological replicates of each strain.

Given that *flg* OFF bacteria in R20291 are phenotypically aflagellate and non-motile (13), we predicted that the six isolates with the flagellar switch in the OFF orientation would be similarly non-motile. The six riboype 012 isolates were examined for the ability to swim through motility medium. Strain 630 and a 630Δ*erm sigD* mutant were used as motile and non-motile controls, respectively (30). After 24 hours incubation, all six ribotype 012 isolates showed a non-motile phenotype in comparison to the motile 630, which migrated outward from the original inoculation site (Figure 5B). Collectively, these data indicate that the flagellar switch is conserved in ribotype 012 clinical and environmental isolates and appear to be “phase-locked” in an *flg* OFF state or undergo flagellar switch inversion at a low frequency *in vitro*.

### Isolation and phenotypic characterization of *flg* ON derivatives of clinical and environmental 012 ribotype isolates

While examining the motility of the ribotype 012 isolates, we observed that a subset of replicates showed motile flares extending from the stab sites for some isolates (Figure 7A). This is in contrast with R20291, for which we previously found that clonal *flg* OFF populations yielded a motile phenotype within 24 hours in 100% of replicates (13). To determine the frequency at which motile flares arise in the ribotype 012 isolates, we increased the number of biological replicates (n = 32 over 2 experiments) and monitored motility every 24 hours for 3 days. A small proportion of colonies of three of the isolates developed motile flares by 24 hours (MT5066, 6.3%; SE838, 9.4%; SS235, 3.1%), and a fourth isolate showed motile flares by 48 hours (MT5065, 18.9 %) (Figure 7B). The number of colonies with motile flares for these four isolates increased and then remained relatively stable between 48 and 72 hours (48 hours: MT5066, 71.9%; SE838, 100%; SS235, 59.4%; MT5065, 18.8%) (Figure 7B). The remaining two isolates, MAM30 and MT5065, failed to develop motile flares throughout the course of the experiments.

**Figure 7.**
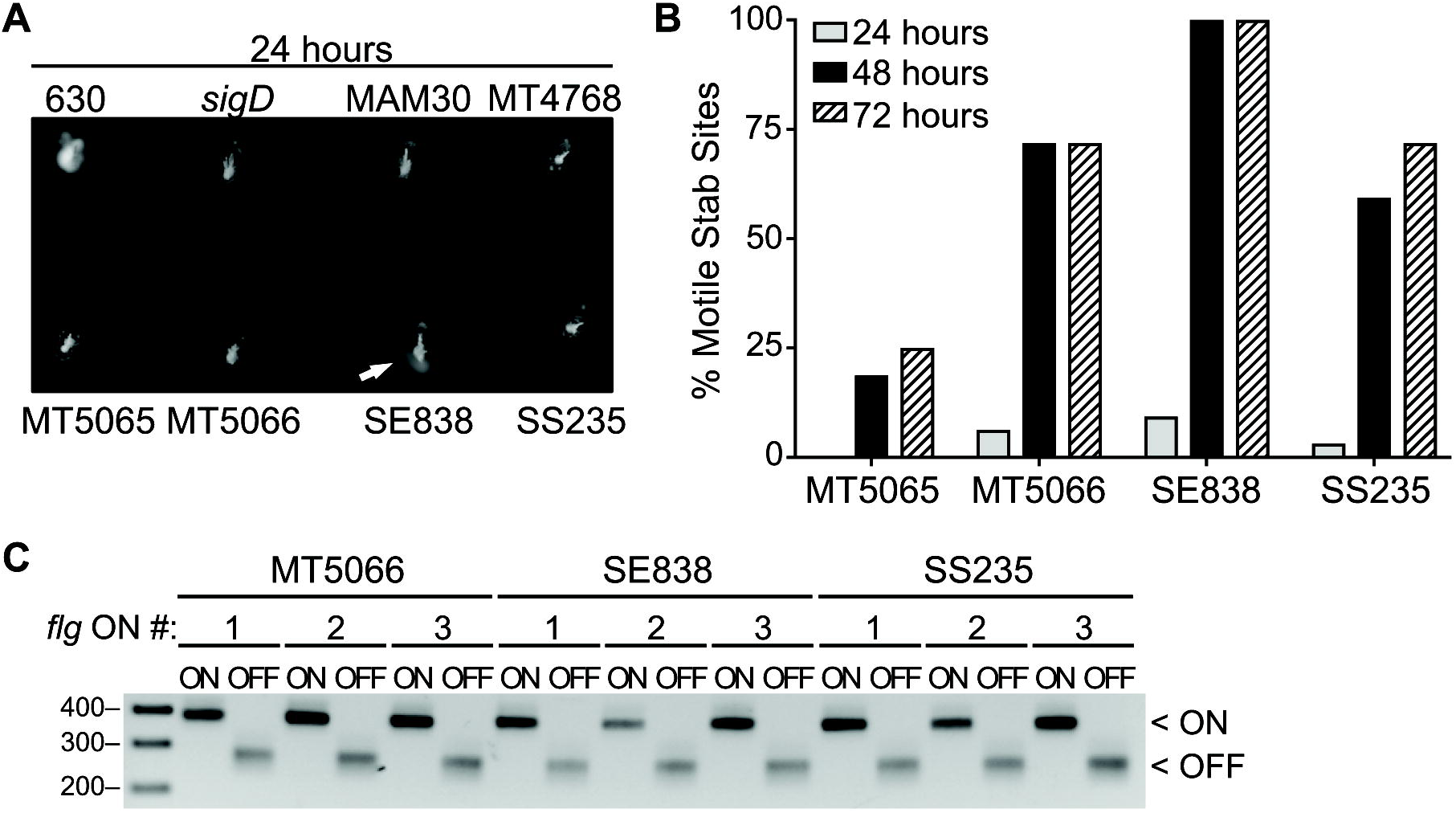
Low frequency recovery of *flg* ON bacteria from select 012 ribotype isolates. (A) Swimming motility after 24 hours. A representative image with a motile flare (white arrow) is shown. (B) The isolates differ in the frequency of recovery of motile revertants. Shown are pooled data from two independent experiments, each examining 16 biological replicates (n = 32). Frequency of motile flares cumulative for each time point, with those detected at 72 hours including those from 48 hours. (C) Orientation-specific PCR products using the indicated strains as template. All nine recovered *flg* ON isolates derived from *flg* OFF MT5066, SE838, and SS235 yielded PCR products indicating both *flg* ON and *flg* OFF populations. Data are representative of three independent experiments.

To confirm that the bacteria from the edge of the flares regained motility through inversion of the flagellar switch to the ON orientation, we arbitrarily chose and isolated bacteria from three motile flares each for MT5066, SE838, and SS235 and subjected them to orientation-specific PCR and sequenced the flagellar switch (Figure 1B). All nine motile isolates yielded a product for the *flg* ON orientation, indicating that DNA inversion occurred in these strains, yet a *flg* OFF product was also observed (Figure S3) (Figure 7C). This result is in contrast to the JIR8094 MD #1-3, for which only the *flg* ON orientation was detected (Figure 2B). In almost all cases, the intensity of the *flg* ON band was greater than the *flg* OFF, suggesting the majority of the population was *flg* ON. Sequencing results confirmed flagellar switch inversion to the ON orientation for all nine *flg* ON derivatives and 100% sequence identity with the strain 630 flagellar switch. We noted that in all three *flg* ON derivatives of parent SE828, the RIR length increased from 22 bp to 23 bp (Figure S3).

To determine whether flagellar switch inversion to the ON orientation was sufficient to restore *flg* ON phenotypes in these strains, we assayed the original ribotype 012 isolates and their *flg* ON derivatives for swimming motility and toxin production (13). As expected, all three *flg* ON derivatives of each strain showed motility compared to their respective non-motile parent strains (MT5066, SE838, and SS235) (Figure 8A). TcdA protein levels were visibly higher in cell lysates of the *flg* ON derivatives relative to the *flg* ON MT5066, SE838, and SS235 strains (Figure 8B). These results demonstrate that the flagellar switch is capable of inversion in some but not all clinical and environmental ribotype 012 isolates, and reiterate the regulatory link between toxin and flagellar gene expression via SigD. In addition, the ribotype 012 strains exhibit heterogeneity in their ability to phase vary, which may be attributed to differences in the sequences of the inverted repeats flanking the flagellar switch.

**Figure 8.**
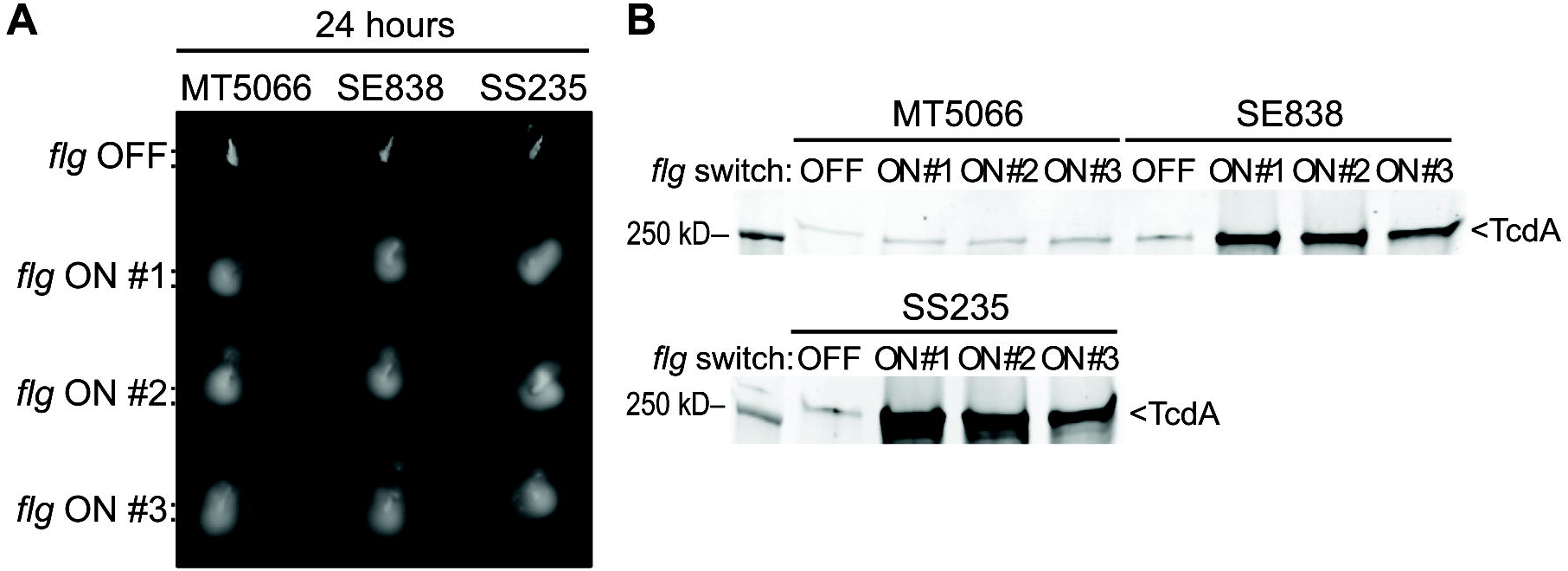
Characterization of select *flg* ON ribotype 012 isolates. (A) Swimming motility assay after 24 hours. All nine recovered *flg* ON isolates were motile at 24 hours compared to their non-motile *flg* OFF MT5066, SE838, and SS235 parents. Data are representative of three independent experiments with five biological replicates for each strain. (B) Detection of TcdA in the indicated strains by western blot. Data are representative of three independent experiments.

## Discussion

Our prior study characterizing flagellum and toxin phase variation in *C. difficile* focused on the epidemic-associated 027 isolate R20291. In this strain, phase variation occurred readily *in vitro*, and the strain consisted of a mixture of *flg* ON and OFF bacteria. However, we noted that the ribotype 012 strain 630, which was originally isolated from a patient with *C. difficile* infection, existed primarily as *flg* ON bacteria. Here, we asked whether the 630 lineage is unusual, or whether reduced phase variation is a broader attribute of ribotype 012 strains. Strain 630’s non-motile JIR8094 derivative primarily exists as *flg* OFF bacteria, and phase variation to *flg* ON occurs with low frequency. Our survey of six clinical and environmental ribotype 012 isolates of *C. difficile* indicates that within the 012 ribotype there exists a range of abilities to phase vary flagellum and toxin production, at least from the *flg* OFF to ON states. This work represents the first examination of flagellar and toxin phase variation within a ribotype family, and our results indicate potentially diverse patterns of phase variation across *C. difficile* strains.

We found that the laboratory-adapted strain JIR8094 contains the flagellar switch in the OFF orientation, which is consistent with its non-motile and toxin-attenuated phenotypes. We were able to isolate rare motile derivatives of JIR8094. These were determined to have inverted the flagellar switch to the ON orientation, suggesting the recovery of motility was attributable to flagellar switch inversion. However, phenotypic characterization of these motile derivatives of JIR8094 indicated that motility and toxin production were only partially recovered compared to the original 630 strain (JIR8094 parent). However, JIR8094 contains 11 polymorphisms compared to 630 (19), and we found that restoring expression of one of the affected genes, *topA*, was sufficient to fully restore motility. This finding indicates that both loss of *topA* and inversion of the flagellar switch from ON to OFF negatively affected flagellar gene expression in JIR8094, and underscores previously stated cautions in using and interpreting results from research done with JIR8094. For example, one study using strain JIR8094 indicated that TcdA is dispensable for virulence in the hamster model (31), while another using 630Δ*erm* showed that both TcdA and TcdB are sufficient for eliciting disease in hamsters (32). The initial studies suggesting TcdA was dispensable for infection may have been due to the *flg* OFF status of JIR8094. A *tcdA* mutation in the background of a motile derivative of JIR8094 could be attenuated for virulence in the hamster model supporting a role for TcdA.

The conditions that favor inversion from *flg* ON to OFF in 630 may be gleaned from how JIR8094 was generated, specifically through serial passage on solid medium (20). In our studies with R20291, we observed a bias toward the *flg* OFF orientation during growth on nutrient rich solid medium over several days, while switch orientation appeared stable during growth in the same liquid medium (13). In contrast to JIR8094, strain 630Δ*erm* was generated by passaging 630 in non-selective liquid medium until a spontaneous erythromycin sensitive isolate was identified, and the flagellar switch remained in the *flg* ON orientation, unlike 630E (21). The ability to bias a phase variable genetic switch in one orientation over another is common in other mucosal bacterial pathogens. For example, incubation in human urine biases the Type I fimbrial switch to the OFF orientation in uropathogenic *Escherichia coli* (33). Anoxic conditions bias the fimbrial switch to the ON orientation in both uropathogenic *Proteus mirabilis* and *E. coli* (34). For *C. difficile*, factors that bias the flagellar switch orientation could be host factors sensed by gene products that influence recombination.

All six of the original ribotype 012 isolates, both clinical and environmental, existed primarily, and perhaps exclusively, in a *flg* OFF state, both genotypically and phenotypically. Two strains, MAM30 and MT4768, appeared to be phase-locked, as we were unable to recover *flg* ON bacteria upon growth in motility medium, which selects for outgrowth of motile bacteria. The remaining four strains varied in their propensity to switch to *flg* ON. One of these, SE838, readily switched from *flg* OFF to ON, similar to R20291. A limitation of our study is that it is uncertain whether the isolates were *flg* OFF at the time of collection, or whether the strains switched during outgrowth in the laboratory. However, the fact that all six isolates from different patients and environments were *flg* OFF argues that our findings are not an artifact of *in vitro* work, but a common feature in the *C. difficile* ribotype 012 family. Recent work found that in a patient simultaneously infected with three *C. difficile* isolates, the ribotype isolate 012 was *flg* ON, providing evidence that not all ribotype 012 isolates are biased to *flg* OFF, or the isolate may have arisen from a stage of infection favoring *flg* ON bacteria (35). It is also possible that most ribotypes show a strong *flg* OFF bias and lower inversion frequency, similar to the ribotype 012, and the ribotype 027 and 017 strains examined previously are outliers (13). A comprehensive analysis of strains representing a variety of ribotypes both *in vitro* and during infection will help determine the population dynamics of flagellum and toxin phase variation in *C. difficile*.

Inverted repeat length may affect recombination of the flagellar switch in the 012 ribotype strains. Our previous analysis of the flagellar switch with publically available sequenced genomes of *C. difficile* indicated 21 bp inverted repeats in all strains. Interestingly, all of the clinical and environmental 012 ribotype isolates had both inverted repeats less than or greater than, but not equal to 21 bp (Figure S1, S2). In addition, the SE838 isolate had flagellar switch IRs of 23 and 22 bp, but its *flg* ON derivatives had a RIR of 23 bp, suggesting some flexibility in the ability of the recombinase RecV to interact with these inverted repeats. It is also possible that a recombination directionality factor (RDF) is required for flagellar switch inversion in *C. difficile* strains that harbor non-ideal inverted repeats, and that the RDF is only produced under certain environmental conditions. The involvement of an RDF may allow precise control over when to promote recombination at the flagellar switch, similar to the contribution of an RDF that contributes to the excision of a DNA element and promotes sporulation (36). Alternatively, changes to repeat length could reduce affinity for RecV, and perhaps allow recombination by another recombinase. Current work seeks to identify a role for inverted repeat sequence and length in affecting flagellar switch inversion frequency in both directions in all ribotypes.

Flagellar and toxin phase variation in *C. difficile* creates population heterogeneity that has the potential to influence diagnosis, disease progression, and transmission. Characterization of flagellar and toxin phase variation in multiple ribotypes of *C. difficile* could provide insight into factors that influence flagellar switch inversion and find correlations between diarrheal disease symptoms and switch orientation. Identifying surface and exported proteins in *C. difficile* that are consistently produced and not subject to phase variable expression could serve as better diagnostic and therapeutic targets to ensure targeting of the entire population.

## Acknowledgments

We thank Victoria Madden from the Microscopy Services Laboratory, UNC-CH Department of Pathology and Laboratory Medicine for guidance with transmission electron microscopy. We thank Robert McKee and Elizabeth Garrett for feedback on this manuscript.

This work was supported by NIH award R01-AI107092 to R.T., and by the UNC-CH Initiative for Maximizing Student Diversity grant from the NIGMS (R25-GM055336), a National Research Service Award Individual Predoctoral Fellowship to Promote Diversity in Health-Related Research grant from the NIAID (F31-AI120613), a UNC-CH Dissertation Completion Fellowship, and a GlaxoSmithKline Science Achievement Award from the United Negro College Fund to B.R.A-F. N.M.V. was supported by a fellowship from the UNC-CH Postbaccalaureate Research Education Program grant from the NIGMS (R25-GM089569). The UNC Microscopy Services Laboratory is supported in part by P30-CA016086 Cancer Center Core Support Grant to the UNC Lineberger Comprehensive Cancer Center from the National Cancer Institute. The funders had no role in study design, data collection and analysis, decision to publish, or preparation of the manuscript.

---

**Supplemental Figure 1. Motile derivatives of JIR8094 arise at a low frequency**. JIR8094 and R20291 cultures were concentrated, and then serial dilutions were assayed for motility in motility medium, with 630 and an R20291 *sigD* mutant as motile and non-motile controls, respectively.

**Supplemental Figure 2. Sequence alignment of flagellar switch and inverted repeat sequences from the clinical and environmental ribotype 012 *C. difficile* isolates**.

**Supplemental Figure 3. Sequence alignment of flagellar switch and inverted repeat sequences from the motile derivatives of clinical and environmental ribotype 012 *C. difficile* isolates**.

**Supplemental Table 1. Strains and plasmids used in this study**.

**Supplemental Table 2. Oligonucleotides used in this study**.

